# Effects of chronic cannabidiol in a mouse model of naturally occurring neuroinflammation, neurodegeneration, and spontaneous seizures

**DOI:** 10.1101/2022.03.07.483344

**Authors:** Joshua T. Dearborn, Hemanth R. Nelvagal, Nicholas R. Rensing, Stephanie M. Hughes, Thomas M Wishart, Jonathan D. Cooper, Michael Wong, Mark S. Sands

## Abstract

Cannabidiol (CBD) has gained attention as a therapeutic agent and is purported to have immunomodulatory, neuroprotective, and anti-seizure effects. Here, we determined the effects of chronic CBD administration in a mouse model of CLN1 disease (*Cln1^-/-^*) that simultaneously exhibits neuroinflammation, neurodegeneration, and spontaneous seizures. Proteomic analysis showed that putative CBD receptors are expressed at similar levels in the brains of *Cln1^-/-^* mice compared to normal animals. *Cln1^-/-^* mice received an oral dose (100mg/kg/day) of CBD for six months and were evaluated for changes in pathological markers of disease and seizures. Chronic cannabidiol administration was well-tolerated, high levels of CBD were detected in the brain, and markers of astrocytosis and microgliosis were reduced. However, CBD had no apparent effect on seizure frequency or neuron survival. These data are consistent with CBD having immunomodulatory effects. It is possible that a higher dose of CBD could also reduce neurodegeneration and seizure frequency.

## 1. Introduction

Cannabidiol (CBD) is a phytocannabinoid found in the *cannabis sativa* plant that has been reported to have anti-seizure, anti-inflammatory, and neuroprotective properties, among a long list of other purported medical benefits^1,2^. When added to an existing anti-seizure drug regimen, chronic CBD is safe and effective at reducing the frequency of seizures in patients with Dravet Syndrome, Lennox-Gastaut Syndrome, and Tuberous Sclerosis Complex^3–6^. In addition, CBD alone exerts acute anti-seizure effects in mouse and rat models of drug-, thermal-, audiogenic-, and kindling-induced seizures^7–11^. However, there are little or no data regarding the effects of CBD in models with concurrent, naturally occurring neuroinflammation, neurodegeneration, and spontaneously arising seizures. In addition, there is a paucity of data regarding the effects of chronic (months) administration of CBD in such murine models of human disease. Most previous studies determined the effects of acute (days to weeks) administration of CBD. We set out to systematically test the effects of chronic, orally-delivered CBD in a well-characterized model that develops all of the above pathologies and clinical signs. The mouse model of CLN1 disease (*Cln1^-/-^*; infantile neuronal ceroid lipofuscinosis; infantile Batten disease) naturally exhibits neuroimmune reactivity, neuroinflammation, neurodegeneration, and spontaneously-arising seizures.

CLN1 disease is a fatal neurodegenerative lysosomal storage disorder that presents with blindness, motor deterioration, cognitive decline, seizures, and early death^12,13^. It results from mutations in the *CLN1* gene, which encodes the soluble lysosomal enzyme palmitoyl protein thioesterase-1 (PPT-1) and is an inherited autosomal recessive disease^14,15^. Palmitoyl protein thioesterase-1 is responsible for removing the fatty acyl chains from palmitoylated proteins. In the absence of PPT-1, undegraded autofluorescent material accumulates in cells throughout the body. However, the most acute and severe disease symptoms are a result of the effects on the central nervous system^12,15^. Currently there is neither a cure nor an FDA-approved therapy for CLN1 disease^13^. Treatment is limited to palliative care, and controlling seizures is challenging and a high priority to caregivers^16^. The goal of this research is to evaluate which aspects of CLN1 disease (neuroimmune response, neurodegeneration, neuroinflammation, and seizures) chronic CBD can impact, and how it may influence disease outcome. We provide empirical evidence of how CBD alone affects these disease characteristics, and add unique data to the relative dearth of research examining how chronic administration of this cannabinoid affects a mouse model of *naturally-occurring* seizures.

## 2. Results

### 2.1 Proteomic analysis

Before embarking on this study, it was critical to determine if putative CBD-binding receptors were present in the brains of *Cln1^-/-^* mice, and if they were present at near WT levels. Therefore, we performed quantitative proteomic profiling of cortical lysates at 7 months of age to confirm that these targets of CBD^2,17–19^ are present in the *Cln1^-/-^* brain. Results from a sampling of CBD-binding partners are shown in Table 1 and reveal that many of the suggested targets of CBD are detected at >85% of levels seen in WT, even at this advanced disease stage.

**Table 1.** Proteomic analysis revealed that many putative CBD targets remain intact in *Cln1^-/-^* mice despite widespread neurodegeneration at 7 months of age.

| Putative CBD targets |  |  |
| --- | --- | --- |
| Gene | Description | *Cln1*<sup>-/-</sup>/WT |
| *Cnr1* | Cannabinoid receptor 1 | 0.959 |
| *Cnrip1* | Cnr-interacting protein 1 | 0.882 |
| *Oprm1* | Mu-type opioid receptor | 0.871 |
| *Slc6a1* | GABA transporter 1 | 0.712 |
| *Slc6a11* | Gaba transporter 3 | 0.933 |
| *Gabarapl1* | GABA receptor-associated protein-like 1 | 0.946 |
| *Gabarapl2* | GABA receptor-associated protein-like 2 | 0.933 |
| *Adora1* | Adenosine receptor A1 | 1.072 |

### 2.2 CBD administration

We set out to determine the effects of chronic CBD treatment on the neuroimmune response, neurodegeneration, and spontaneous seizures in *Cln1^-/-^* mice. Our initial attempt at dosing *Cln1^-/-^* mice with CBD involved multiple daily intraperitoneal (i.p.) injections. The half-life of CBD is much shorter in mice (~4 h) compared to humans (18-32 h)^11,17,20,21^, resulting in the necessity for multiple daily administrations. However, as *Cln1^-/-^* mice age, they become overreactive to handling and will often display seizure and/or spastic activity when handled. Mice that were treated via *i.p*. injection, whether with CBD or vehicle, also had a shorter lifespan compared to untreated/unhandled mice (data not shown) which led us to believe that repeated handling and injection alters the disease phenotype in an undesirable manner. As such, we shifted to a voluntary oral administration method. Mice were administered CBD via flavored gelatin cube 5 days per week beginning at PND 30 and continuing through PND 209.

It was critical to confirm delivery of the pure CBD to the brain through this oral route. Cannabidiol was quantified in the brains of three CBD- and vehicle-treated mice. The amount of CBD was below the level of quantification (LOQ) in the three vehicle-treated mice. In contrast, the three CBD-treated mice had 61.99 ng/g, 221.18 ng/g, and 71.08 ng/g of CBD in brain tissue (Table 2). This confirms that voluntary oral consumption of CBD via gelatin cubes delivers measurable levels of drug to the brain.

**Table 2.** Voluntary oral administration via gelatin cube delivers quantifiable levels of CBD to the brain.

|  | Cannabidiol |  |
| --- | --- | --- |
|  | Mean (ng/g) | SD |
| *Cln1*<sup>-/-</sup> + CBD | 118.1 | 89.4 |
| *Cln1*<sup>-/-</sup> + Vehicle | <LOQ | <LOQ |

Limit of quantification (LOQ) was 1.5ng/g tissue

### 2.3 Neuroimmune and neuroinflammatory responses

CD68 and GFAP staining are increased in untreated *Cln1^-/-^* mice, and these markers of glial activation are commonly used as a proxy for a neuroimmune or neuroinflammatory response. Chronic CBD treatment reduced CD68 expression in S1BF (*t*(8) = 3.949, *p* = 0.0042) as well as GFAP expression in VPM/VPL of the thalamus (*t*(8) = 2.651, *p* = .0292) (Fig. 1a-h). We previously showed that the levels of cytokines and chemokines are altered in untreated *Cln1^-/-^* mice compared to WT mice^22–24^. Specifically, pro-inflammatory cytokines such as TNF-α and IFN-γ are typically increased in the mutant mice, and certain treatments are successful at reducing them^25^. We analyzed cytokines/chemokines in treated and untreated *Cln1^-/-^* mice to survey for anti-inflammatory effects. The markers analyzed can be grossly divided into three categories: anti-inflammatory cytokines (IL-10, IL-4, and IL-5), monocyte activators (IP-10, MCP-1, and MIP-1β), and pro-inflammatory cytokines (IL-1β, IL-6, IFNγ/, IL-12p70, GRO-α, and TNF-α). In brain tissue exposed to CBD for 6 months, none of the markers were significantly altered when compared to tissue exposed to vehicle only (*p* ≥ 0.146 for all; Fig. 1i).

**Figure 1.**
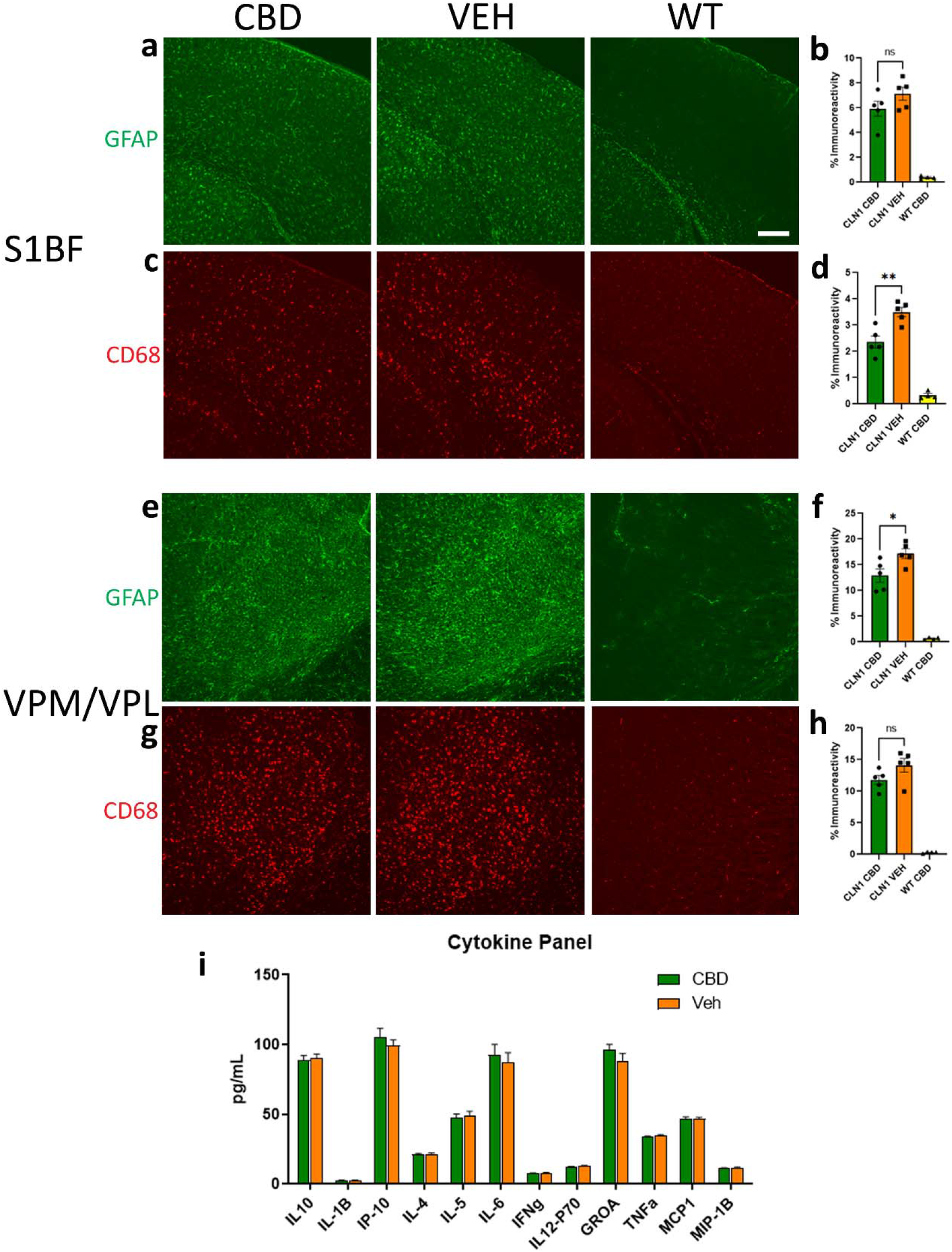
Treatment with CBD reduces some disease-related pathology. In S1BF, chronic CBD did not affect GFAP immunoreactivity (a,b) but did significantly reduce CD68 expression (c,d). Conversely, GFAP expression was reduced following CBD treatment (e,f) while CD68 immunoreactivity was unchanged (g,h) in VPM/VPL. Chronic CBD did not significantly affect any of the cytokines analyzed (i). Data shown are means ± SEM. *Indicates *p* < 0.0292; **indicates *p* = 0.0042 Scale bar is 200 μm.

### 2.4 Cellular/Morphological markers of disease

Untreated *Cln1^-/-^* mice consistently display cortical atrophy. Although some treatments that target the underlying enzymatic defect can preserve cortical thickness^26,27^, there was no significant increase in cortical thickness in the CBD-treated animals (Fig. 2a, b). In addition, chronic CBD treatment did not affect neuron count in either the cortex or the thalamus in *Cln1^-/-^* mice (Fig. 2c, d). Finally, SCMAS is a major component of the undegraded storage material in CLN1 disease. Cannabidiol did not reduce this histological marker of disease (not shown).

**Figure 2.**
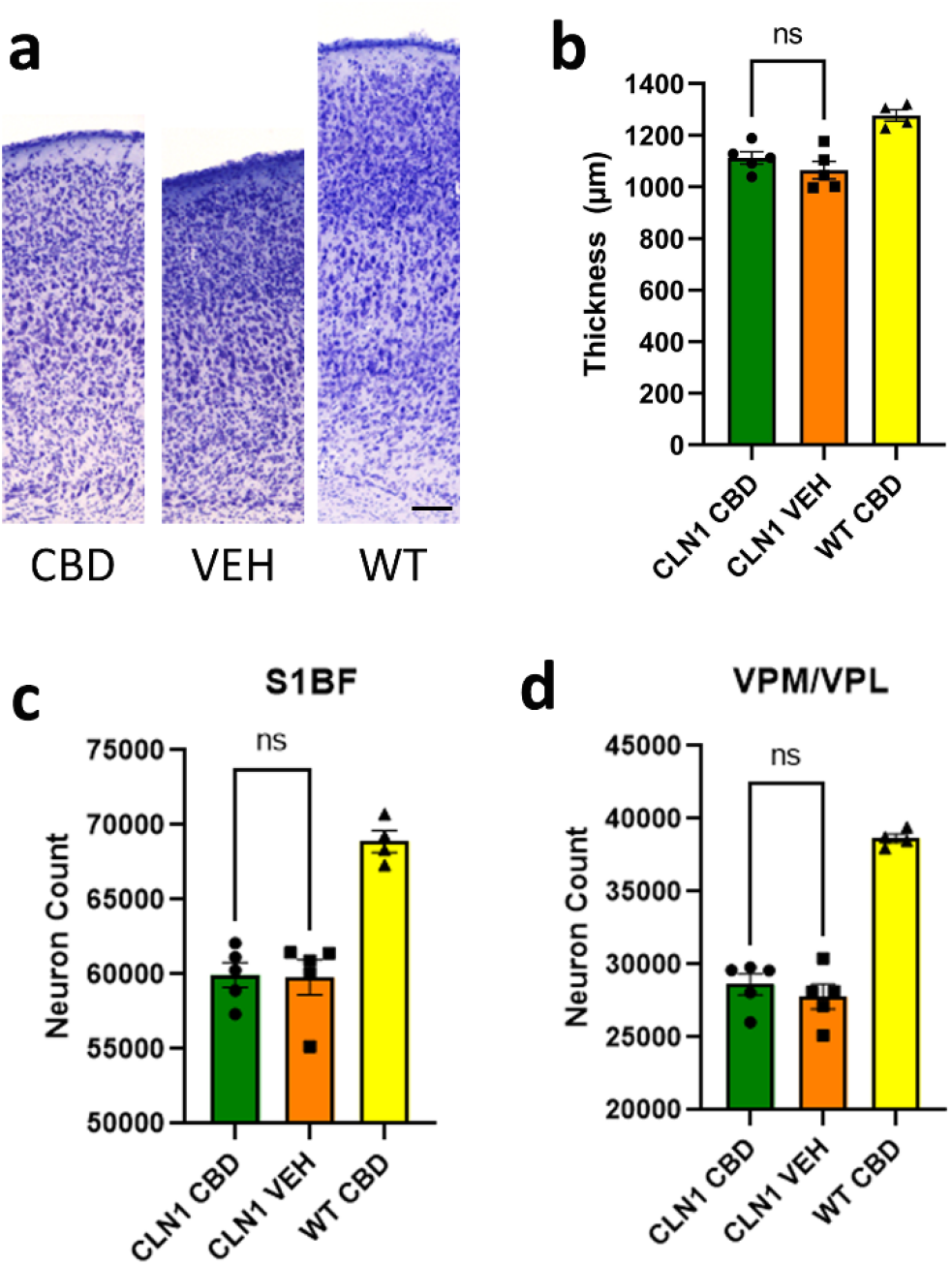
Chronic CBD does not affect neuron count or cortical thickness. (a) Nissl-stained depiction of cortical thickness. Treatment did not affect cortical thickness (b) or neuron count in S1BF cortex (c) or VPM/VPL of the thalamus (d). Scale bar is 250 μm.

### 2.5 Seizure trajectory

We previously showed that *Cln1^-/-^* mice display seizures beginning at approximately 7 months of age; specifically, 50% of *Cln1^-/-^* mice display seizures (defined as electrographic discharges longer than 10 sec) at that age^28,29^. However, these data were based on relatively small numbers of mice (n = 7 or fewer and n = 9 or fewer, respectively^28,29^) for a relatively short duration of EEG monitoring (48 consecutive hours). Before committing to treating mice chronically prior to seizure monitoring, we conducted a more comprehensive evaluation of the development and trajectory of seizures in the *Cln1^-/-^* mouse for the purpose of identifying an ideal window for EEG monitoring. Simultaneous video and EEG monitoring demonstrated concordance between epileptiform discharge and behavioral, “popcorn-like” seizures (Supplemental Video S1). In the current evaluation, we monitored for seizures in a much larger cohort (n = 20) of mice for a considerably longer period of time than previously reported. Continuous EEG-monitoring in *Cln1^-/-^* mice began on PND 180 (approximately 6 months of age), and more than 1 mouse exhibited at least 1 seizure on that first day. For evaluation across time, seizure data was batched into 5-day bins. Day-to-day variability was extremely large at times (from 0 seizures to 55 seizures on any given day), and seizures were often clustered, therefore bins provided more normalized data that allowed for evaluation of seizures across many days. The mean number of seizures across mice increased steadily until approximately PND 215, with the first death of a mouse occurring at PND 209 (Fig. 3). By PND 190, 70% of *Cln1^-/-^* mice exhibited at least 1 seizure per 5-day bin. Between PND 200 and 210, that proportion rose to 80%. Prior to PND 190, fewer than 60% of mice exhibited at least 1 seizure per bin, and after PND 216 the mean number of seizures decreases until the last mice alive exhibit fewer than 1 seizure per 5-day bin (Fig. 3). Based on these data, we determined that the optimal observation window for monitoring seizures in *Cln1^-/-^* mice is between PND 190 and 209. Within that timeframe, all of the monitored mice were still alive, and 70% or more exhibited at least 1 seizure per bin.

**Figure 3.**
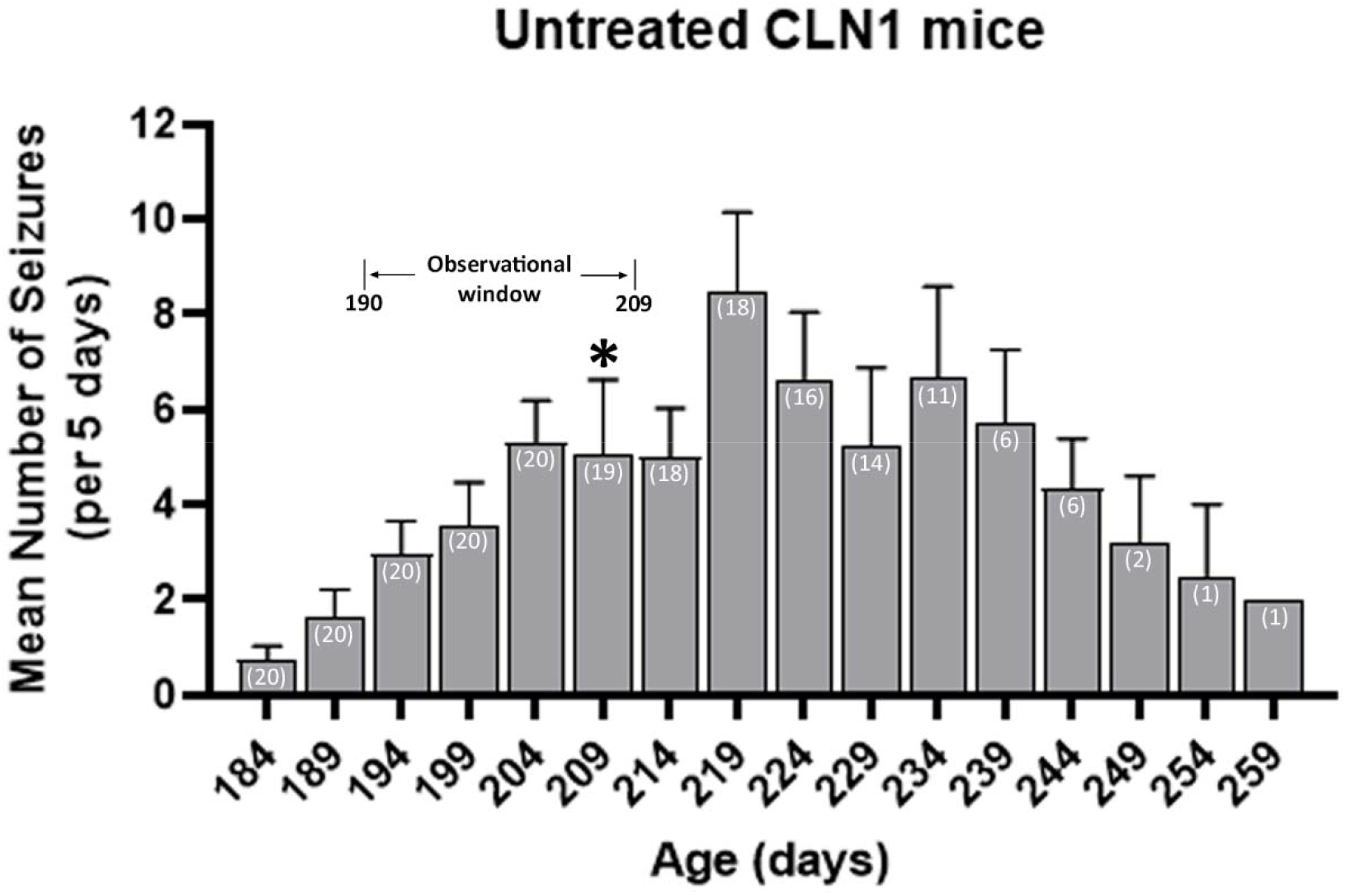
Trajectory of seizures in *Cln1^-/-^* mice. Seizure occurrences were summed into 5-day batches, and mean seizure counts (+SEM) are shown. The first of 20 mice died at PND209, indicated with an asterisk. Number of surviving mice is included in white text within each bar. Selected observational window is shown here, indicating the span during which seizures were measured in the cohort of mice receiving chronic treatment. Data is shown for all living mice until the last mouse died at PND 259.

There was no difference in life span between untreated, EEG-monitored males and females (Fig. 4a). However, male and female *Cln1^-/-^* mice differ when it comes to seizure frequency. Females exhibit more total seizures compared to males during the timespan when all mice remained alive [PND 180 to PND 209, *t*(18) = 2.246, *p* = 0.0375], and they appear to have an earlier onset (Fig. 4b). During this time period, a higher percentage of female mice exhibited at least 1 seizure earlier compared to males (Fig. 4c). Finally, more females exhibited seizures across this timeframe than males as measured by area under the curve [AUC; *t*(18) = 3.513, *p* = 0.0025] (Fig. 4d).

**Figure 4.**
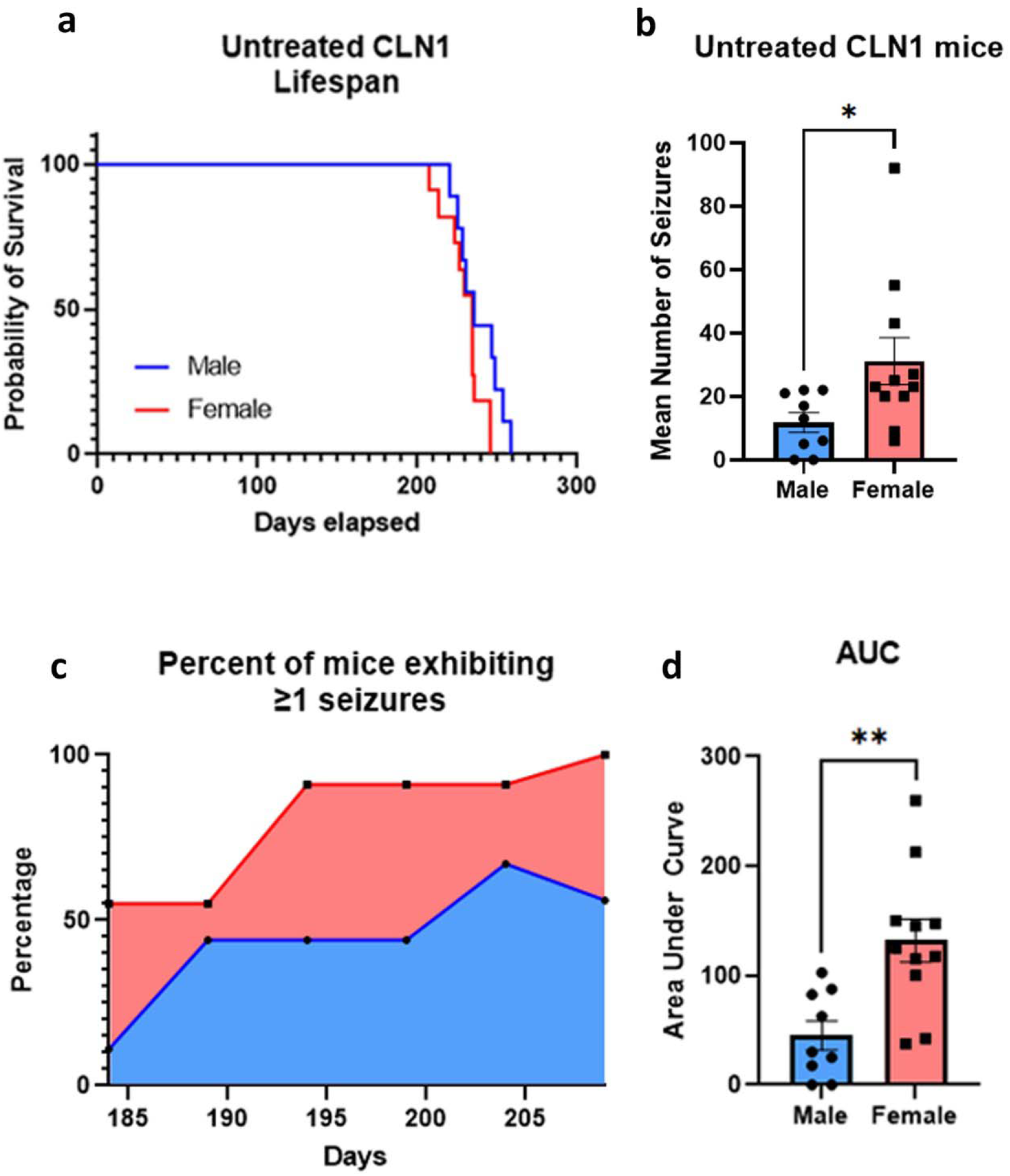
Female *Cln1^-/-^* mice exhibit more, and earlier, seizures, compared to males. (a) While the four longest-lived mice were all male, there was no difference in life span between untreated, EEG-monitored males (n=9) and females (n=11). (b) Females exhibited more total seizures, over the period when all animals remained alive, compared to males, and they appeared to have an earlier onset. (c,d) A higher percentage of female mice exhibited at least 1 seizure earlier compared to males, and more females exhibited seizures across this timeframe than males as measured by area under the curve. Data shown are means ± SEM. *Indicates *p* = 0.038; **indicates *p* < 0.0025

### 2.6 Chronic CBD and seizures

Chronic treatment with CBD does not appear to qualitatively affect the EEG pattern of epileptiform discharge associated with behavioral seizures in *Cln1^-/-^* mice, nor does it alter total seizure frequency(Fig. 5a,b). There were also no differential effects of CBD on seizures based on sex (Fig. 5c). Compared to *Cln1^-/-^* mice receiving treatment with vehicle (mean = 19.61, SD = 11.79), mice receiving treatment with CBD (mean = 20.11, SD = 16.90) experienced a similar number of seizures during the observation window [*t*(35) = 0.1026, *p* = 0.9189] (Fig. 5d, e).

**Figure 5.**
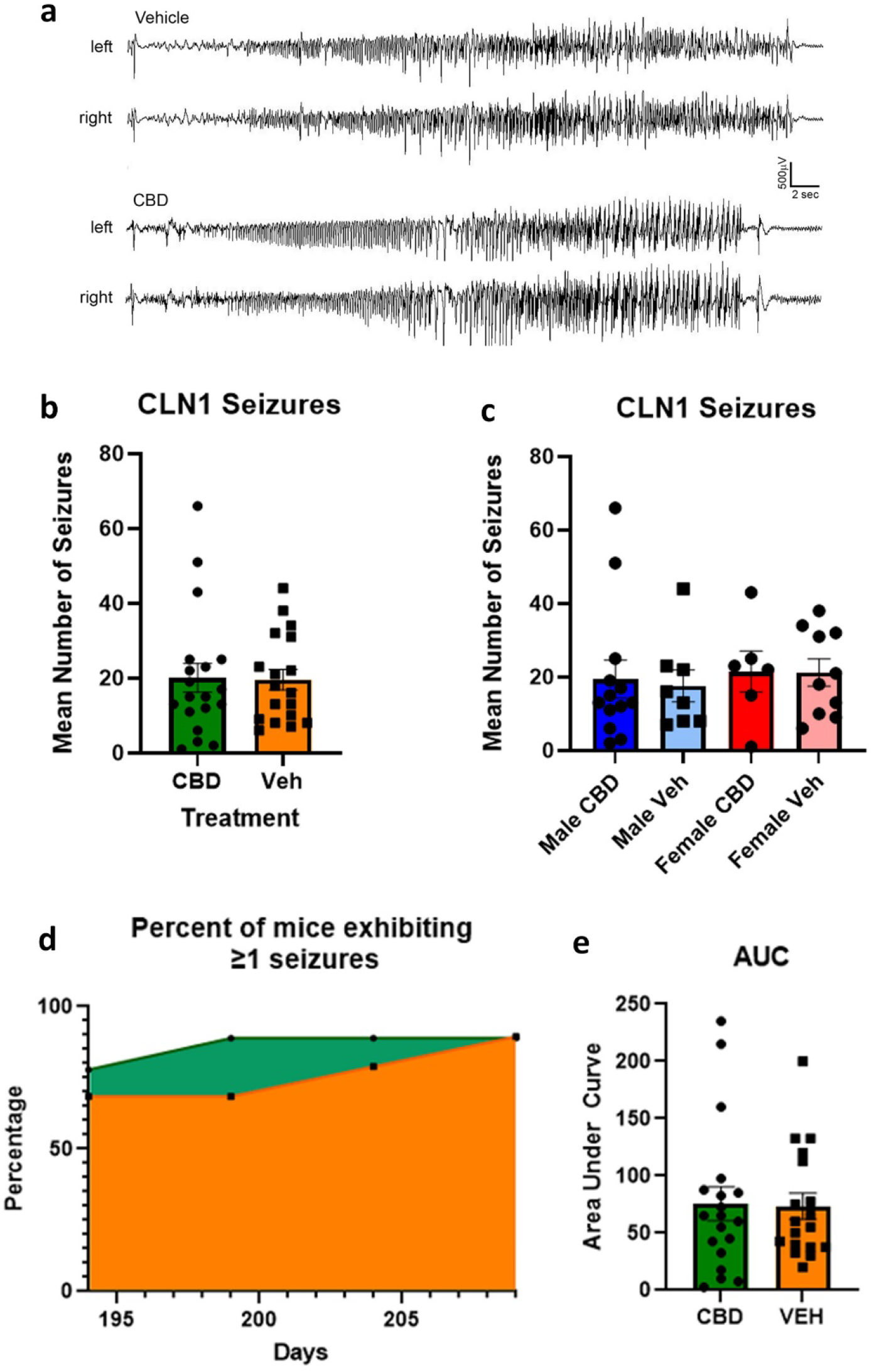
Chronic treatment with cannabidiol does not affect seizures in *Cln1^-/-^* mice. (a) EEG tracings during seizures are displayed for *Cln1^-/-^* mice treated with either vehicle or CBD; readings from electrodes placed on the left and right side of the skull are shown. Mice treated with CBD or vehicle exhibited similar numbers of seizures, and this was true within both males and females when analyzed separately (b,c). A similar proportion of *Cln1^-/-^* mice exhibited seizures across the observation window regardless of treatment, as shown in the area under the curve analysis (d,e). Data shown are means ± SEM.

## 3. Discussion

The goal of this research was to determine the effects of chronic, orally-administered cannabidiol on a variety of phenotypes, including neuroinflammation, neurodegeneration, and spontaneous seizures. The CLN1 mouse model reliably and naturally displays all of these pathological and clinical signs. It is important to note that we used highly purified cannabidiol, not a “whole-plant” extract. Whole-plant extracts typically contain hundreds, if not thousands, of chemicals native to the *cannabis sativa* plant. We sought to elucidate the individual contribution of the increasingly-popular and widely-available cannabinoid, CBD.

Our LC-MS/MS proteomic data confirmed that putative targets of CBD are present at near normal levels in the *Cln1^-/-^* mouse brain, even near the end stage of disease. This is interesting in light of the fact that CLN1 disease is profoundly neurodegenerative and suggests that neurons expressing CBD receptors may be partially and preferentially spared. This also provided the impetus to proceed with the CBD study. Importantly, chronic oral administration of ~100mg/kg of CBD was well tolerated and had no obvious deleterious effects. Not unexpectedly, chronic administration of CBD had no effect on the progressive accumulation of SCMAS in the brains of treated *Cln1^-/-^* mice. Cannabidiol is not known to mediate any mechanism that would increase PPT1 enzyme activity or otherwise decrease SCMAS accumulation. Therefore, it is important to keep in mind that any effects of CBD would be observed in the context of continued disease progression.

Markers of a neuroimmune response, such as astrocytosis (GFAP) and microgliosis (CD68), are dramatically increased in the untreated *Cln1^-/-^* mouse brain and can be observed as early as 1-3 months of age^22,24,28^. Here we show a significant decrease in both GFAP and CD68 in areas of both the thalamus and the cortex following chronic CBD treatment. This reduction of cellular markers of disease is remarkable considering the continued disease progression and the fact that the animals were evaluated at a relatively late stage of disease. A growing body of literature reports anti-inflammatory effects of CBD^1,2^. In the current study, whole brain homogenates from animals treated with CBD showed neither a decrease in pro-inflammatory nor an increase in anti-inflammatory cytokines/chemokines. It is possible that chronic treatment with CBD might have had a more pronounced effect earlier in the disease process. The apparent discordance between the cellular and humoral responses could be due to the different analytical approaches. Immunofluorescent imaging allows for the sensitive detection of regional differences. In contrast, regional differences in cytokine/chemokine levels could be masked when analyzing whole brain homogenates. We chose to analyze cytokine levels at the whole brain level because 1) receptors targeted by CBD are spread throughout the brain, 2) neuropathology is widespread by late disease stage in CLN1, and 3) the etiology of seizures in the *Cln1^-/-^* mouse is unknown, so probing only specific brain regions may have been limiting. Alternatively, CBD could have differential effects on the cellular and humoral responses through as yet unknown mechanisms. Future studies will be required to determine the effects of CBD treatment at various time points to determine the temporal nature of the reductions in astrogliosis and microglial activation. In addition, it will be critical to determine if the effects of CBD are enhanced when combined with treatments that target the underlying defect in CLN1 disease, PPT1 deficiency. It is possible that CBD could synergize with another therapy to reduce both the cellular and humoral immune responses.

One of the hallmarks of CLN1 disease is profound neurodegeneration and cortical atrophy. In fact, at a terminal stage of disease, the mass of an affected child’s brain can be as little as 50% that of an age-matched normal child^14,15^. Likewise, the *Cln1^-/-^* mouse also has profound neurodegeneration, cortical atrophy and significantly decreased brain mass^26,30^. Although there appeared to be a slight increase in cortical thickness with CBD treatment, this did not reach statistical significance. In addition, there was no significant change in neuron counts in either the thalamus or the cortex following chronic CBD treatment. Similar to the neuroimmune response, it will be important to determine the temporal effects on neurodegeneration and whether or not CBD can synergize with therapies designed to increase PPT1 activity.

Here we report a more comprehensive characterization of the spontaneous epilepsy phenotype in the mouse model of INCL. From our data it is clear that *Cln1^-/-^* mice experience seizures earlier than previously reported^28,29^. Seizures clearly develop prior to 7 months of age, and are likely to appear, albeit at a low frequency, even before 6 months of age. This finding tells us what recent pathological data^22,28,31,32^ have revealed in abundance: treatment is most likely to be successful when initiated early in the disease process. This supports our strategy of beginning CBD treatment at 1 month of age, which is prior to the development of any measurable pathological or clinical signs. Our findings also uncovered a significant sex difference in seizure frequency in untreated *Cln1^-/-^* mice, with female mice exhibiting significantly more seizures than males during the optimal observation period. A higher proportion of female mice exhibit at least one seizure per day at an earlier time point than male mice, a difference that persists through the end of life. This is consistent with previous findings in CLN3 disease patients (juvenile Batten disease) where females typically display a more severe trajectory of symptoms compared to males^33^. Our findings also align with research in the CLN6 (variant late infantile NCL) mouse in that onset of some behavioral symptoms were earlier and more severe in females compared to males^34,35^. Although it is tempting to extrapolate findings between various forms of CLN disease, this should be viewed cautiously since these diseases can have fundamentally different causes. For example, both CLN3 and CLN6 disease result from defects in transmembrane proteins, whereas CLN1 involves deficiency in a soluble lysosomal enzyme. Regardless, the current findings reveal the merit of further research into the nature of sex differences in CLN1 disease.

It has been shown previously that CBD protects against seizures in multiple mouse models of induced epilepsy. These include models of PTZ-, audiogenic-, and thermal-induced seizures^7–11^. We are unaware of the use of CBD to reduce naturally occurring seizures in a murine model. In addition, the overwhelming majority of studies administer CBD over a relatively short period of time (days to weeks). The current study differs significantly from previous work in that the *Cln1^-/-^* mouse naturally develops seizures as a result of increasing disease pathology (i.e., there is no need to artificially induce seizures). Given that CLN1 disease is progressive, and the spontaneous seizures appear later in the disease, we also administered CBD for at least six months. It is clear that chronic, oral administration of CBD alone does not significantly reduce seizure frequency. In addition, treatment with CBD did not delay the onset of seizures in the *Cln1^-/-^* mouse. Given the reduction in astrocytosis and microgliosis, the complete lack of any effects on seizures is perhaps a bit surprising. We previously showed that decreasing these cellular markers in the *Cln1^-/-^* mouse via a small molecule, brain-penetrant anti-inflammatory drug (Minozac) is associated with a small but significant decrease in seizure frequency^25^. One explanation for this apparent discrepancy is that Minozac was specifically designed as an antiinflammatory agent and is likely more potent than CBD.

Anecdotally, caregivers of patients with infantile Batten disease report anti-seizure effects of *cannabis* products. This could be due to what is commonly referred to as the “entourage effect”, which hypothesizes that phytocannabinoids work best in the context of terpenes and flavonoids that occur naturally in the *cannabis sativa* plant. It is possible that a “whole-plant” *cannabis* product would provide greater benefit to pathological markers of disease, or seizure frequency, compared to CBD alone. Unfortunately, due to the relatively crude nature of such products it is virtually impossible to determine the specific effects of CBD, even if it is one of the major components of these preparations. Further, as is the case with many human conditions that involve epilepsy, patients are often taking a regimen of 3-5 antiseizure drugs prior to the addition of a *cannabis* product^3–5^. This makes it even more difficult to determine any positive effects attributable solely to CBD or any other phytocannabinoid. It is certainly possible that CBD could enhance the effects of other anti-seizure drugs or vice versa. It has been reported that CBD can increase the concentration of active metabolites of other antiseizure medications. The most robust evidence comes from CBD-clobazam interactions^36^.

Where previous research has shown various protective effects of CBD at lower doses than used here^1,37–40^, the lack of an effect on seizure occurrence reported in the murine model of CLN1 disease may be explained by some key differences. First, route of administration affects brain and plasma concentrations of CBD greatly; *i.p*. delivery leads to a faster and greater concentration of drug in the brain compared to oral delivery^17^. As noted above, repeated *i.p*. delivery negatively affects *Cln1^-/-^* mice, as would repeated oral gavage, so the voluntary oral dosing was the most practical and clinically relevant route of administration. Still, it is clear that we delivered measurable drug to the brain, which peaks at 6 h following oral administration^17^. Another important consideration is the use of induced states of epilepsy/inflammation/pain in the literature compared to the spontaneous seizures that develop in *Cln1^-/-^* mice. This murine model displays profound neurodegeneration that progresses rapidly, leading to naturally-occurring seizures that may simply be more difficult to treat compared to seizure models that must be induced via external intervention. Finally, we tested only one dose of CBD in the current study; evaluation of various doses used in previous work^1,38,39^ may yield different results.

Taken together, we provide evidence for positive effects of chronic CBD on the neuroimmune response in the mouse model of CLN1 disease. Importantly, this was observed using CBD in the presence of ongoing disease and in the absence of any adjunct antiinflammatory, neuroprotective, or anti-epileptic treatments. Although 100mg/kg of CBD alone is not sufficient to prevent or reduce the naturally occurring seizures in the *Cln1^-/-^* mouse, it might enhance the effects of other anti-seizure drugs or treatment strategies designed to target the fundamental genetic defect. Finally, it is possible that a higher dose of CBD may be necessary to achieve positive anti-seizure benefits. The dose (~100mg/kg) of CBD administered in the current study is within the range reported in the literature(5-600mg/kg)^13,23–25^, but it is possible that an even higher dose is required to have an effect on seizures in the context of such profound disease. Given the thorough characterization of the CLN1^-/-^ mouse seizure phenotype and the ability to chronically and non-invasively administer CBD, these questions can now be addressed directly.

## 4. Materials and Methods

### 4.1 Animals

One cohort of mice consisted of 14 mice [5 *Cln1^-/-^* mice treated with CBD, 5 *Cln1^-/-^* mice treated with vehicle, and 4 untreated wild-type (WT) mice] to be used for immunofluorescence and cytokine analysis. A second cohort of untreated *Cln1^-/-^* mice was used to evaluate the development and trajectory of seizures and determine an optimum seizure observation window; this cohort consisted of 20 mice (9 males, 11 females). A final cohort of mice was used to evaluate the effects of chronic CBD treatment on seizures during the observation window. This cohort consisted of a total of 37 *Cln1^-/-^* mice, with 19 (13 males) receiving CBD treatment for 6 months and 18 (8 males) receiving treatment with vehicle for 6 months. From this cohort of 37 mice, brains from 3 CBD-treated and 3 vehicle-treated mice were used to determine the level of CBD by mass spectrometry in the brain following chronic exposure. In all cohorts, mice were single-housed from postnatal day (PND) 30 to control for access to treatment. *Cln1^-/-^* mice were first generated by Gupta *et al^30^* and maintained on a C57Bl/6J background^41^. Animals were housed in a controlled-access animal facility at Washington University School of Medicine (St. Louis, MO) under a 12 h light/dark cycle. Food and water were available *ad libitum,* and procedures were carried out in accordance with NIH guidelines and under a protocol approved by the Institutional Animal Care and Use Committee (IACUC) at Washington University School of Medicine. This study is reported in accordance with ARRIVE guidelines^42^

### 4.2 Proteomic analysis

Brains from 7-month-old WT and *Cln1^-/-^* mice were prepared for protein extraction and liquid chromatography—tandem mass spectrometry (LC-MS/MS) as detailed in our previous work^43^. Briefly, cortical protein levels were determined using the bicinchoninic acid assay (BCA) (Pierce, UK) and samples (n=4 per genotype) were pooled by group. Tandem mass tags were applied (https://doi.org/10.7488/ds/2750 (2020)) before peptides were fractionated for LC-MS/MS. Raw MS data were searched against mouse (Mus musculus) protein sequences from UniProtKB/Swiss-Prot using the MASCOT search engine (Matrix Science, Version 2.2) through Proteome Discoverer software (Version 1.4, Thermo Fisher). Identifications were quantified as the ratio of expression in *Cln1^-/-^* cortex to that in WT cortex.

### 4.3 Chronic treatment

We chose a moderate to high dose (~100mg/kg) based on the literature evaluating effects of CBD on acutely induced seizures in mice. Such research uses doses that range from as low as 5 mg/kg to as high as 600 mg/kg^10,44–46^. We determined that WT and *Cln1^-/-^* mice eat an average of 60-75% (by weight) of a 1cm^3^ cube of flavored gelatin over a 24 h period and will do so reliably over long periods of time (data not shown). Accordingly, we suspended CBD in ethanol and mixed it with flavored gelatin during preparation. Briefly, raspberry flavored gelatin (Jell-O brand) was prepared as instructed on the package with the addition of extra unflavored gelatin (Knox brand) such that each mL (resulting in a solid 1cm^3^ cube) contained 180 mg of raspberry Jell-O and 140 mg of unflavored gelatin. Cannabidiol (NIDA Drug Supply Program, 99.5% pure) was suspended in 95% ethanol at 400 mg/mL and added such that each resulting gelatin cube contained 4.5 mg of CBD. Because mice reliably eat approximately 2/3 of the cube over the span of 24 h, this would result in an effective dose of approximately 100 mg/kg per 30 g mouse. Mice were given a raspberry-flavored gelatin cube containing either CBD or vehicle (ethanol only) once per day in the afternoon, just before the dark cycle commenced. The following day at the same time, the remnants of that cube were removed and replaced with a fresh cube. This occurred 5 days per week beginning at 1 month of age and ending after the completion of EEG monitoring on PND 210 (approximately 7 months) for a total chronic treatment period of 6 months. To measure seizure activity in the chronically treated mice, surgery, electrode placement, and monitoring procedures were carried out precisely as described below, with the exception that surgery took place on PND 183, monitoring began on PND 190, and the study ended on PND 210.

### 4.4 Tissue collection

For mass spectrometry, chronically treated mice that were subject to EEG-monitoring were removed from the apparatus on PND 210 and euthanized via i.p. injection with Fatal-Plus (Vortech Pharmaceuticals, Dearborn, MI, U.S.A). Mice were perfused with PBS before removal of the brain which was immediately flash-frozen in liquid nitrogen. For histological evaluation of disease markers as well as cytokine expression analysis, chronically treated mice were euthanized via Fatal-Plus, perfused with PBS, and brains removed with one hemisphere flash-frozen in liquid nitrogen and the remaining hemisphere placed in 4% paraformaldehyde in PBS for 48 hours. After 48 h, brain hemispheres were removed from paraformaldehyde and placed in 30% sucrose in 50 mM TBS (pH=7.6) until processing.

### 4.5 Mass spectrometry

To confirm that voluntary oral consumption delivers CBD to the brain, mass spectrometry was performed on 6 mouse brains (3 *Cln1^-/-^* brains treated with CBD and 3 *Cln1^-/-^* brains treated with vehicle). For each mouse, one hemisphere of brain was homogenized with a 5mm Zircon bead, 600μL of acetone, and 15μL of 100ng/mL d3-CBD (Qiagen TissueLyser II (30 hz, 9 min)). The homogenate was sonicated for 10 min, then centrifuged (10,000 rpm, 3 min, 4°C) before the supernatant was removed and transferred to a new tube. Sonication and centrifugation were repeated, and the extractions were combined and dried. Samples were reconstituted in 150μL. MeOH and mixed via vortex and sonication, filtered via VivaClear Mini 0.8 μm, PES spin filters (Sartorius), then transferred to autosampler vials with 175μL inserts for analysis.

Each sample was analyzed in triplicate using LC-MS as follows. 3μL injections of each sample were separated on a C18 column eluted at 15 μL/min with ddH_2_O with 0.1% formic acid (A) and acetonitrile with 0.1% formic acid (B) using the following gradient: 30% B (0-2 min), 30% B to 100% B (2-18 min), 100% B (18-24 min), and reequilibrated at 30% B (24-30 min). Sample ionization utilized a heated ESI source with the following settings: spray voltage +3600 v, sheath gas 27 (arb), auxiliary gas 0.4 (arb), sweep gas 0 (arb), ion transfer tube 325 °C, and vaporizer temperature ambient. CBD and d3-CBD (IS) were detected using SRM (selected reaction monitoring). For CBD, the precursor ion was 315.2 m/z, and 259.21 m/z (collision energy 28.38 V) and 193.16 m/z (collision energy 32.34 V) were the product ions. For d3-CBD, the precursor ion was 318.3 m/z, and 262.21 m/z (collision energy 29.35 V) and 196.20 (collision energy 34.32 V) were the product ions. The SRM properties were as follows: cycle time 0.8 s, Q1 resolution 0.7 FWHM, Q3 resolution 1.2 FWHM, CID gas (N2) 1.5 mTorr, and source fragmentation 0.

### 4.6 Immunostaining

A one-in-six series of coronal brain sections from each mouse were stained on slides using a modified immunofluorescence protocol^22^ for the following antibodies - astrocytes (rabbit anti-GFAP, 1:1000, Agilent Z0334), microglia (rat anti-mouse CD68, 1:400, Bio-Rad MCA1957), and subunit c of mitochondrial ATPase (SCMAS) (rabbit anti-SCMAS, 1:200, Abcam ab181243). Coronal sections were cut at 40 μm on a Microm HM430 freezing microtome (Microm International GmbH, Wallendorf, Germany). The sections were mounted on Superfrost plus slides (Fisher Scientific) and air-dried for 30 min, slides were then blocked in a 15% serum solution (Normal goat serum, S-1000 Vector Laboratories) in 2% TBS-T (1x Tris Buffered Saline, pH 7.6 with 2% Triton-X100, Fisher Scientific) for 1 h. Slides were then incubated in primary antibody in 10% serum solution in 2% TBS-T for 2 h. Slides were washed three times in 1xTBS and incubated in fluorescent Alexa-Fluor-labelled IgG secondary antibodies (Alexa-Fluor goat anti-rabbit 488 Invitrogen A-11008, goat anti-rat 546 Invitrogen A-11081 all 1:400 dilution) in 10% serum solution in 2% TBS-T for 2 h, washed three times in 1xTBS and incubated in a 1x solution of TrueBlack lipofuscin autofluoresence quencher (Biotium, Fremont, CA) in 70% ethanol for 2 min before rinsing in 1xTBS. Slides were coverslipped in fluoromount-G mounting medium with DAPI (Southern Biotech, Birmingham, AL).

### 4.7 Cresyl fast violet staining

A one-in-six series of 40 μm coronal sections were stained for cresyl fast violet (Nissl) as before^31,32^, to reveal their cytoarchitecture. Briefly, brains were mounted on chrome-gelatin coated Superfrost microscope slides (ThermoFisher Scientific, MA, USA) and air-dried overnight. All sections were incubated for 60 min at 60°C in 0.1% cresyl fast violet and 0.05% acetic acid. Stained sections were then differentiated through a series of graded ethanol solutions (70%, 80%, 90%, 95% and 2×100%) (Fisher Scientific, MA, USA) before clearing in Xylene (Fisher Scientific, MA, USA) and coverslipping with DPX (Fisher Scientific, MA, USA), a xylene-based mountant.

### 4.8 Measurements of cortical thickness

Cortical thickness measurements were obtained blinded to genotype and treatment, from cresyl fast violet stained slides, as previously described^31^, using StereoInvestigator software (MBF Bioscience Inc, Williston, VT). Contours were drawn over the primary somatosensory barrel cortex (S1BF), making 10 measurements per section over 3 consecutive sections in accordance to defined anatomical landmarks^47^ at 5x magnification.

### 4.9 Thresholding image analysis

To analyze the degree of glial activation in the gray matter (GFAP-positive astrocytes + CD68-positive microglia) as well as accumulation of storage material (SCMAS), a semiautomated thresholding image analysis method was used with Image-Pro Premier software (Media Cybernetics)^22^. Briefly, this involved the collection of slide-scanned images at 10x magnification for a one-in-six series of sections per animal followed by demarcation of all regions of interest^47^ while maintaining the lamp intensity, video camera setup, and calibration constant throughout image capturing. Images were subsequently analyzed using Image-Pro Premier (Media Cybernetics) using an appropriate threshold that selected the foreground immunoreactivity above background. This threshold was then applied as a constant to all subsequent images analyzed per batch of animals and reagent used to determine the specific area of immunoreactivity for each antigen^22^. Specific regions of interest included the ventral posteromedial/posterolateral (VPM/VPL) nuclei of the thalamus and the S1BF cortex^28,31,32^.

### 4.10 Stereological Analysis

Estimates of neuron populations in the Ventral posteromedial/ lateral nuclei (VPM/VPL) of the thalamus and the barrel field of the primary somatosensory cortex (S1BF) were performed using a design-based optical fractionator method in a one-in-six series of Cresyl fast violet stained sections (West et al., 1991 Anat Rec 231:482-497) using Stereo Investigator software (MBF Bioscience). Cells were sampled with counting frames (90 ×100 μm) distributed over a sampling grid (VPM/VPL, 350 × 350 μm; S1BF, 450 × 450 μm) that was superimposed over the region of interest at 100× magnification.

### 4.11 Cytokine assays

Quantification of chemokines and cytokines in the brain was performed as previously described^23^. Briefly, a 12-biomarker Multi-Analyte Profile was generated for *Cln1^-/-^* mice by MYRIAD RBM using standard Luminex technology. The analytes included the following: IL-10, IL-1β, IP-10, IL-4, IL-5, IL-6, IFN-γ, IL-12p70, GRO-α, TNF-α, MCP-1, and MIP-1β. Following euthanasia and tissue removal, brains were homogenized in a solution of 10 mM Tris (pH 7.5), 150 mM NaCL, 0.1% Triton X-100, and 20 μL/mL protease inhibitor cocktail (Sigma). The supernatant was diluted to a protein concentration of approximately 0.5-1.0 mg/mL and stored at −80°C until analysis. Cytokine concentrations were quantified using standard MYRIAD RBM protocols.

### 4.12 Electroencephalography (EEG) electrode surgery

*Cln1^-/-^* mice received surgery for placement of EEG electrodes at PND 173 (or PND 183 in for treated mice). Custom wire EEG electrode sets were constructed using five Teflon coated stainless steel wires (76 μm bare diameter) soldered to an electronic pin header and a micro screw. The soldered contacts were covered with dental cement and the electrode set sterilized for implantation.

Mice were placed under 3-4% isoflurane on a stereotaxic frame with a heating pad set to 36.5°C until pedal withdraw reflex ceased. The skin was prepared with betadine and alcohol wipes with isoflurane maintained at 1-1.5% for the remainder of the procedure. After a midline vertical incision to expose the skull, forceps and 3% hydrogen peroxide were used to remove any connective tissue and dry the skull for electrode placement. Burr holes for the frontal reference electrodes were made (anterior +0.5 mm, lateral +/-0.5 mm; bregma) using a micro drill with a 0.9 mm tip and screws were secured in the skull. Two bilateral “active” recording electrodes were placed over the parietal cortex (posterior −2.5 mm, lateral +/- 1.5; bregma) and a ground screw secured over the cerebellum (posterior −6.2 mm, lateral +/- 0.5; bregma) using the same technique as the reference electrode. The exposed skull, screws, and all wires were covered in a layer of dental cement (SNAP, Parkell) with the pin header secured to the head for subsequent recording. The skin was sutured around the exposed dental cement/pin header and tissue glue (Vetbond, 3M) used to close the remainder of the incision. Mice received Buprenorphine (0.1 mg/kg) and recovered in a warmed chamber for two hours then placed in individual recording cages^48^.

### 4.13 Video-EEG monitoring and baseline seizure frequency

Groups of four mice (up to 16 at a time) in individual caging recovered from surgery for 72 h prior to connection with a custom flexible cable attached to the exposed pin header for recording. Bilateral cortical EEG signals were acquired using a referential montage using Stellate acquisition software and amplifiers. Signals were amplified at 10,000X with high-pass (0.5Hz) and low-pass (100Hz) filters applied. EEG signals were digitized at 250Hz and time-locked video EEG collected continuously^48^. EEG signal reflective of seizure activity was matched against corresponding video recording to confirm the behavioral phenotype, an often violent episode wherein the mouse stiffens into a rearing posture and is propelled about the chamber in a “popcorn-like” manner. Twenty *Cln1^-/-^* mice (9 males, 11 females) were continuously monitored for EEG activity starting at PND 180 until the animals died or were sacrificed for humane reasons.

### 4.14 Statistical analysis

Analyses were performed using IBM SPSS Statistics version 27 and GraphPad Prism 9 using two-tailed unpaired *t* test when assumptions of normality are met, and Kruskal-Wallis test when assumptions of normality are violated. A *p* value < 0.05 was considered statistically significant. Data are expressed as mean ± standard deviation (SD) except where indicated otherwise.

## Acknowledgments

Funding for this research was provided by:

Batten Disease Support and Research Association to JTD
NIH NS R01100779 to MSS
NIH P50 HD103525 to MW
NIH NS R21 106523 to MSS
BBSRC BBS/E/D/10002071 to TMW

## Author contributions

JTD, MW, and MSS designed the project. JTD, HRN, NRR, and TMW performed the research. JTD, NRR, and HRN analyzed the data. SMH, TMW, and JDC contributed procedures and reagents. JTD, HRN, NRR, and MSS wrote the manuscript.

